# Conditional deletion of *Ccl2* in smooth muscle cells does not reduce early atherosclerosis in mice

**DOI:** 10.1101/2023.08.06.552177

**Authors:** Stine Gunnersen, Jeong Tangkjær Shim, Fan Liu, Uwe J.F. Tietge, Charlotte Brandt Sørensen, Jacob Fog Bentzon

## Abstract

**Background and aims:** C-C motif chemokine ligand 2 (CCL2) is a pro-inflammatory chemokine important for monocyte recruitment to the arterial wall and atherosclerotic plaques. Global knockout of *Ccl2* reduces plaque formation and macrophage content in mice, but the importance of different plaque cell types in mediating this effect has not been resolved. Smooth muscle cells (SMCs) can adopt a potentially pro-inflammatory function with expression of CCL2. The present study aimed to test the hypothesis that SMC-secreted CCL2 is involved in early atherogenesis in mice.

**Methods:** SMC-restricted Cre recombinase was activated at 6 weeks of age in mice with homozygous floxed or wildtype *Ccl2* alleles. Separate experiments in mice lacking the Cre recombinase transgene were conducted to control for genetic background effects. Hypercholesterolemia and atherosclerosis were induced by a tail vein injection of recombinant adeno-associated virus (rAAV) encoding proprotein convertase subtilisin /kexin type 9 (PCSK9) and a high-fat diet for 12 weeks.

**Results:** Unexpectedly, mice with SMC-specific *Ccl2* deletion developed higher levels of plasma cholesterol and larger atherosclerotic plaques with more macrophages compared with wild-type littermates. When total cholesterol levels were incorporated into the statistical analysis, none of the effects on plaque development between groups remained significant. Importantly, changes in plasma cholesterol and atherosclerosis remained in mice lacking Cre recombinase indicating that they were not caused by SMC-specific CCL2 deletion but by effects of the floxed allele or passenger genes.

**Conclusions:** SMC-specific deficiency of *Ccl2* does not significantly affect plaque development in hypercholesterolemic mice.

**Bullet points:**

- SMCs express CCL2 in human plaques and upon inflammatory activation of a murine SMC line in vitro.
- Deletion of the Ccl2 gene in SMCs using Cre recombinase does not influence the size or composition of atherosclerotic plaque in mice.
- Only by conducting a control study in mice without Cre recombinase was an initially observed difference in atherosclerosis concluded to be a genetic background effect.

## Introduction

Atherosclerosis is a low-grade, non-resolving inflammatory disease driven by retention of low-density lipoproteins (LDL) in the arterial wall [1]. Inhibition of inflammatory pathways reduces plaque progression in experimental models and protects against the incidence of clinical atherosclerotic events in patients with prior myocardial infarction [2,3]. Blocking inflammation broadly in the body, however, adversely affects defenses against infection. Research is needed to identify mechanisms that locally control inflammatory cells to devise potent anti-inflammatory therapies that are specific or selective for atherosclerosis.

C-C motif chemokine ligand 2 (CCL2) is a pro-inflammatory chemokine secreted by multiple cell types, which is central in the recruitment of monocytes to tissues and has a clear causal role in atherosclerosis [4,5]. Global deficiency of *Ccl2* in *Apoe*^-/-^ mice, *Ldlr*^-/-^ mice, and mice overexpressing apoB consistently inhibited atherogenesis and reduced plaque content of macrophages [6–8]. Blocking *Ccl2* activity in mice with advanced atherosclerosis reduced further plaque progression and changed plaque composition to contain fewer macrophages, more SMCs, and more collagen [9]. These studies indicate a continued role of CCL2 that extends beyond disease initiation. CCL2 is also expressed in human atherosclerotic plaques and associated with features of plaque vulnerability [10], and *CCL2* polymorphisms and elevated CCL2 plasma levels are associated with an increased risk of cardiovascular disease [10–12].

Expression in myeloid cells accounts for a part of the atherosclerosis-promoting effect of CCL2 in mice [13], but local arterial cells also express CCL2 and may carry some of the proatherogenic effect [4,14,15]. Several studies have suggested roles for modulated SMCs in controlling plaque inflammation. Cultured SMCs can be induced by interleukin-1β (IL-1β) and minimal modified low-density lipoprotein particles (mmLDL) to express CCL2 and other pro-inflammatory genes [16,17]. In *Ldlr*^-/-^ mice, SMC-specific deficiency of inhibitor of nuclear factor kappa B kinase subunit beta (IKKβ), a catalytic subunit necessary for activation of nuclear factor kappa B (NF-κB) that is upstream of *Ccl2* expression, had multiple metabolic effects, including adipose tissue changes and reduced plaque development [18]. Furthermore, SMC-specific conditional knockout of the interleukin-1 receptor (IL-1R) substantially reduced plaque formation in *Apoe^-/-^* mice [19]. Recently, a lineage-tracing study in mice suggested dichotomous roles of SMC-derived CCL2 in atherosclerosis development depending on the phenotypical modulation state of the SMCs [20]. These studies indicate the presence of important effector functions of modulated SMCs for controlling plaque inflammation and progression, but the nature of these is only sparsely understood. In the present study, we assessed the importance of SMC-secreted CCL2 for early atherosclerosis in hypercholesterolemic mice.

## Materials and methods

Please see an expanded methods section in the Supplemental Material.

### Mice

Male *Myh11-*CreER^T2^ mice (B6.FVB-Tg(Myh11-cre/ERT2)1Soff/J, stock no. 019079) expressing tamoxifen-inducible Cre recombinase (*CRE-ERT2)* under the SMC-specific myosin heavy chain 11 (*Myh11*) promotor [21], and *Ccl2*^flx/flx^ mice (B6N.129S1(FVB)-Ccl2tm1.2Tyos/J, stock no. 023347) with *loxP* sites flanking exon 1-2 of the *Ccl2* gene [22] were acquired from Jackson and intercrossed. Experimental mice of our study were male offspring of breeders heterozygous for the floxed *Ccl2* allele. Only males were included because the Myh11-CreER^T2^ transgene is inserted on the Y chromosome. One of the male breeders was found without the Y-linked *Myh11*-*CreER^T2^*, presumably due to the exchange in genetic material between pseudoautosomal regions of X and Y chromosomes [23,24], and was used to breed CreER^T2^-negative male mice for control experiments. Animals were housed in groups with unlimited access to tap water and a standard laboratory diet in an enriched environment with standard bedding and nesting material under a 12/12 h day-night cycle in a temperature (20-25°C) controlled facility.

At 6-8 weeks of age, male *Myh11-*CreER^T2^ x *Ccl2*^flx/flx^ and *Myh11-*CreER^T2^ x *Ccl2*^wt/wt^ littermates were injected i.p. with 1 mg tamoxifen (T5648, Sigma-Aldrich) dissolved in corn oil (P2144, Sigma-Aldrich) daily for 10 days to obtain mice with or without SMC-specific *Ccl2* deletion (*Ccl2*^SMC-KO^ and *Ccl2*^SMC-WT^). Male *Ccl2*^flx/flx^ and *Ccl2*^wt/wt^ littermates were treated similarly. At 12 weeks of age, hypercholesterolemia was induced by a single tail vein injection of rAAV8-D377Y-mPCSK9 virus particles (rAAV-PCSK9, 1 × 10^11^ vector genomes, produced at Vector Core at the University of North Carolina, Chapel Hill, North Carolina, USA) and shifted to a high-fat diet (D12079B, Research Diets Inc.) containing 21% fat and 0.2% cholesterol [25]. After 12 weeks of feeding atherosclerosis-promoting diet, the mice were anesthetized with 5 mg pentobarbital and 4 mg lidocain i.p. and terminated by exsanguination. The arterial tree was perfused with Cardioplex for 30 seconds and 4% phosphate-buffered formaldehyde (PFA) for 5 minutes at 100 mmHg through the left ventricle, followed by immersion-fixation in 4% PFA for 24 hours. Animal procedures were approved by The Danish Animal Experiments Inspectorate (license 2015-15-0201-00542) and conducted at Aarhus University following ARRIVE guidelines.

### Atherosclerosis analysis

The top half of the heart containing the aortic root was removed and embedded in paraffin. Transversal sections (3 μm) of the aortic root were cut from the distal part of the heart and serial sections were collected at 0 μm, 80 μm, 160 μm and 240 μm from the level of the aortic valve commissures. For quantification of atherosclerosis, sections from all levels (main experiment) or the first two levels (control experiment with Cre-negative mice) were stained by orcein. Sections from the first level (main experiment) were stained for smooth muscle alpha-2 actin (ACTA2) and galectin 3 (LGALS3) using mouse monoclonal anti-ACTA2 (Dako, cat. no. M0851, 1:100) after blocking with Fab fragment (Jackson ImmunoResearch, cat. no. 715-007-003, 1:10) and rat monoclonal IgG2a anti-LGALS3 (Cederlane, cat. no. CL8942AP, 1:250) followed by Alexa 488-conjugated donkey anti-mouse (Invitrogen, cat. no. A21202, 1:400) or Alexa 488-conjugated donkey anti-rat (Invitrogen, cat. no. A21208, 1:400) secondary antibodies. Analysis of stained sections was performed in Fiji (also known as ImageJ) [26].

Aortae were cleaned of all periadventitial fat, cut open longitudinally, stained with Oil Red O for 10 minutes at 37^°^C, washed in isopropanol and PBS, and mounted on a microscope slide with Aquatex for en face quantification of lesion coverage in the aortic arch down to the supreme intercostal artery. Analysis of the scanned slides was performed in Fiji.

### Human atherosclerotic plaques

Human atherosclerotic plaques from the left anterior descending artery obtained from a previously published material [23] were used for immunohistochemical analysis of CCL2 localization using rabbit polyclonal anti-CCL2 (Lifespan Bioscience, cat. no. LS-B10540, 1:50) and Envision+ System-HRP (DAB) (Dako, cat. no. K4010). The original material was collected after approval by the regional ethical committee [27]. It was subsequently anonymized and did not require further ethical approval for the use in the present study.

### Cell studies

An immortalized murine smooth muscle cell line (MOVAS, ATCC CRL-2797) was maintained in culture in DMEM (Fischer Scientific, cat. no. 41965062) supplemented with 10% fetal calf serum (FCS) (Sigma-Aldrich, cat. no. F7524), 1% penicillin-streptomycin (Invitrogen, cat. no. 15140122) and 1% gentamycin (G418, Invivogen cat. no. ant-gn-1). Primary human aortic SMCs (hVSMCs), obtained as previously described [24], were kindly provided by Thomas Ledet, Aarhus University. They were maintained in DMEM supplemented with 10% FCS and 1% penicillin-streptomycin. Incubation was performed at 37^°^C in a humidified atmosphere of 95% air and 5% CO_2_ and the medium was changed every 2-3 days. MOVAS cells were seeded for experiments in 12-well plates at a density of 6.25−10^3^ cells/cm^2^, and human SMCs were seeded in 6-well plates with a density of 1.67#x2212;10^4^ cells/cm^2^. The experiments were performed 24 h after seeding the cells. To induce an inflammatory response in hVSMCs and MOVAS cells, the cells were incubated with IL-1β (R&D Diagnostics, cat. no. 401-ML-005/CF, 0.01 – 100 ng/ul) for 24 hours. The conditioned media (CM) were then collected and cleared from debris by centrifugation (3,000xg for 5 min). CCL2 levels were measured in conditioned media by ELISA (Invitrogen, cat nos. 88-7391-88 and 88-7399-88).

### Statistics

All statistical analysis was performed using GraphPad Prism 9 (GraphPad Software Inc.). Data was checked for normal and log-nomal distribution and equal variance and analyzed using the tests indicated in figure legends. Multiple regression analysis was used to analyze associations with plaque data using genotype and plasma cholesterol as independent variables and associations with plasma cholesterol levels (using genotype and plasma PCSK9 concentrations as independent variables). *P* values below 0.05 were defined as statistically significant. Data are shown as mean ± standard error of the mean (SEM) as indicated in each figure legend. The number of animals/samples analyzed is stated in figure legends. In some analyses, data points were not obtained for all mice because of technical failures. All procedures and analyses were performed blinded for the mouse genotype.

## Results

### CCL2 is expressed in human atherosclerotic lesions

To evaluate the expression pattern of CCL2 during human atherogenesis, we performed CCL2 immunohistochemistry in human coronary lesions of different severity (**Fig. 1A**). CCL2 was found expressed in normal human coronary artery in both the medial and intimal layer. In early lesions, CCL2 was present in foam cells, but also in the underlying artery wall. In advanced atherosclerotic lesions, CCL2 was detected in areas adjacent to the necrotic core, while CCL2 expression in the medial layer was diminished. Staining of adjacent sections with ACTA2 showed considerable overlap with the pattern of CCL2 staining suggesting widespread SMC expression of CCL2 in normal media/intima as well as in developing plaques.

**Figure 1.**
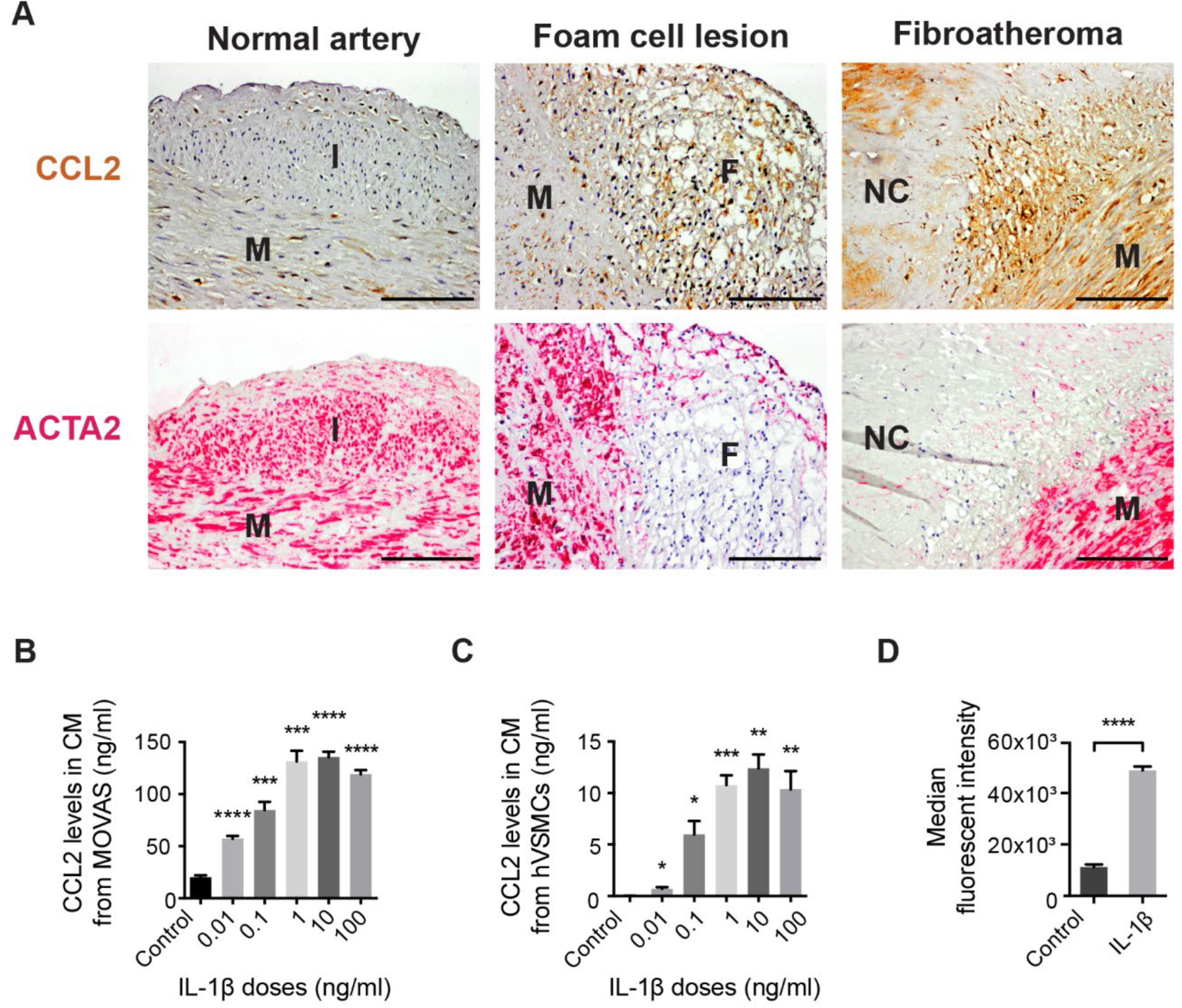
CCL2 expression in SMC-rich regions of human plaques and CCL2 secretion in SMC cultures. (A) CCL2 (brown color) and ACTA2 (red color) detected by immunohistochemistry in adjacent sections of normal human artery wall and atherosclerotic lesions of different severity. CCL2 is expressed in media and intima of the normal arterial wall. CCL2 is also expressed in foam cells and underlying medial SMCs, and in active inflammatory areas adjacent to the necrotic core in the fibroatheroma. F, foam cells. I, Intima. M, Media, NC, necrotic core. Scale bars = 100 μm. CCL2 is secreted from cultured SMCs stimulated with IL-1β. CCL2 levels measured in conditioned media (CM) from (B) MOVAS cells (n=6 replicates per group) and (C) hVSMCs (n=5-6 replicates per group) treated with IL-1β for 24 hours. (D) Flow cytometry measurement of intracellular CCL2 accumulation in MOVAS cells after blocking the secretory pathway by Brefeldin-A. IL-1β stimulation (1 ng/ml) increases CCL2 accumulation measured as median fluorescent intensity (n=3 replicates per group). Graphs show mean ± SEM. \**p*<0.05, \*\**p*<0.01, \*\*\**p*<0.001, \*\*\*\**p*<0.0001 by Dunnett’s T3 multiple comparisons test following Brown-Forsythe ANOVA (A,B) and unpaired Student’s t-test (D).

### Human and murine SMCs in culture can be stimulated to express CCL2

Previous studies have reported CCL2 secretion from murine SMCs upon stimulation [28]. To confirm this, we stimulated human VSMCs and a murine SMC line (MOVAS) with IL-1β and found that the secretion of CCL2 in the conditioned media of both increased dose-dependently (**Fig. 1B-C**). Also, assessed by flow cytometry, we found an increase in protein expression of CCL2 in response to IL-1β stimulation of MOVAS cells (**Fig. 1D**).

### SMC-specific conditional knockout of *Ccl2* in hypercholesterolemic mice

Modulation to a CCL2-secreting SMC phenotype by local IL-1β or other pro-inflammatory stimuli could facilitate ongoing inflammation in the developing atherosclerotic lesion. To test this hypothesis *in vivo*, we crossed mice with SMC-restricted CreER^T2^ expression and either homozygous floxed or wildtype *Ccl2* alleles to obtain mice with SMC-specific deletion of *Ccl2* (*Ccl2*^SMC-KO^) and littermate controls (*Ccl2*^SMC-WT^) (**Fig. 2A**). Mean allelic recombination efficiency was 75.9 ± 1.3% (mean ± SEM) (**Fig. 2B**). Because recombination was determined in aortic samples after mechanical removal of the adventitia and endothelium, the analyzed cells may have contained a small amount of non-SMC cell types causing some underestimation of recombination efficiency.

**Figure 2.**
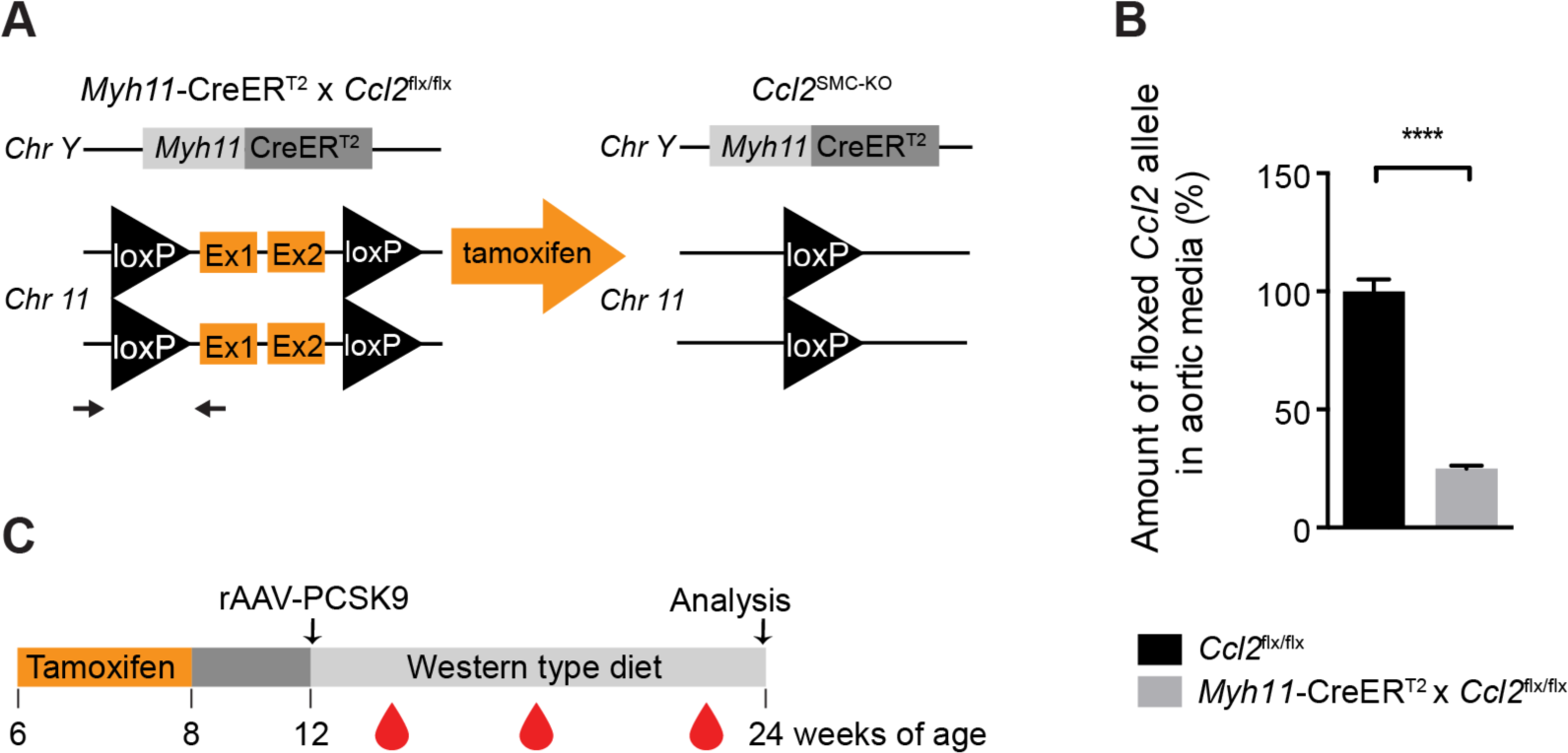
Experimental design. (A) Illustration of the genetic alterations in *Myh11-*CreER^T2^ x *Ccl2*^flx/flx^ mice that lead to SMC-specific knockout of *Ccl2* after tamoxifen injection. Black arrows flanking the left loxP site indicate primers used for detecting recombination by qPCR. (B) Recombination efficiency was measured as the reduction in (non-recombined) floxed CCL2 alleles in genomic DNA of the aortic media in *Myh11-*CreER^T2^ x *Ccl2*^flx/flx^ (n=21, grey bar) compared with *Ccl2*^flx/flx^ (n=15, black bar) mice. The graph shows mean ± SEM. \*\*\*\**p*<0.0001 by unpaired Welch t-test. (C) Schematic illustration of the mouse experiment showing the timing of tamoxifen injections (10 daily doses between 6-8 weeks of age), induction of LDL receptor-deficiency by a single tail-vein injection of rAAV8-D377Y-mPCSK9 at 12 weeks of age, and final analysis after feeding Western-type diet for 12 weeks. Blood samples were collected 2, 6, and 11 weeks after rAAV-PCSK9 injection.

### Increased plasma cholesterol levels in *Ccl2*^SMC-KO^ mice

Atherosclerosis was induced in *Ccl2*^SMC-KO^ and *Ccl2*^SMC-WT^ mice by rAAV-PCSK9 injection followed by feeding a high-fat diet for 12 weeks (**Fig. 2C**). Plasma cholesterol was similar in the two genotypes after 2 weeks but significantly higher in *Ccl2*^SMC-KO^ mice than *Ccl2*^SMC-WT^ mice at both 6 and 11 weeks (**Fig. 3A**). Analysis of size-fractioned lipoproteins measured at 12 weeks showed that the increased cholesterol in *Ccl2*^SMC-KO^ mice was in apoB-containing lipoproteins (VLDL and LDL) (**Fig. 3B**).

**Figure 3.**
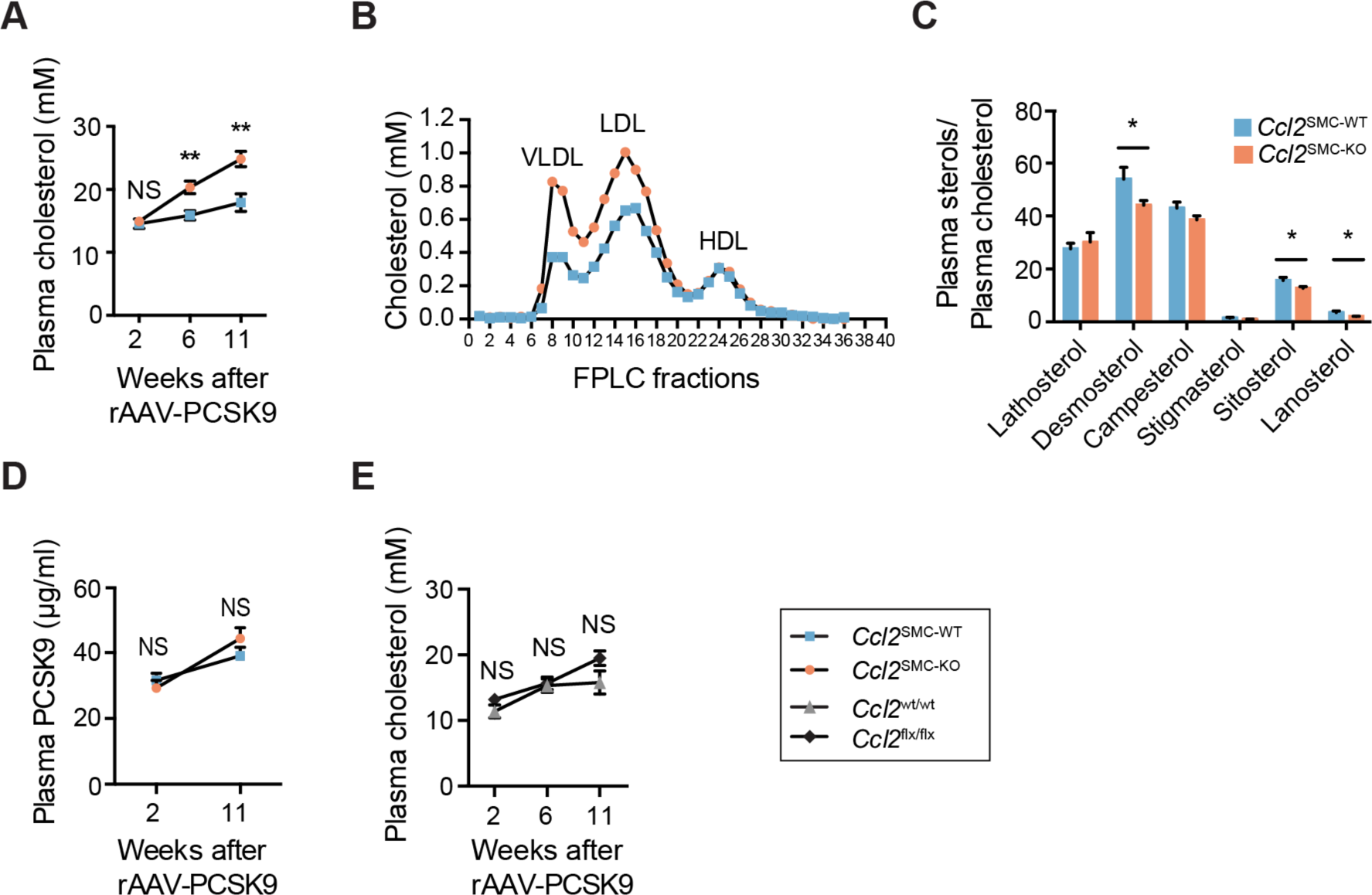
Plasma cholesterol phenotype. (A) Plasma cholesterol levels were initially similar but significantly increased in *Ccl2*^SMC-KO^ (n=22, orange circles) compared with *Ccl2*^SMC-WT^ (n=24, blue squares) mice at 6 and 11 weeks after rAAV-PCSK9 injection. (B) Analysis of size-fractionated lipoproteins by FPLC at 12 weeks after rAAV-PCSK9 injection revealed increases in cholesterol in VLDL- and LDL-sized, but not HDL-sized, lipoproteins. (B) Sterol analysis for markers of cholesterol biosynthesis (lathosterol and desmosterol) or cholesterol absorption (campesterol, stigmasterol, sitosterol, lanosterol) showed no significant changes or changes opposite the direction that could explain the increased plasma cholesterol levels in *Ccl2*^SMC-KO^ (n=23) compared with *Ccl2*^SMC-WT^ mice (n=21). Plasma PCSK9 levels were not different between *Ccl2*^SMC-WT^ (n=22) and *Ccl2*^SMC-KO^ mice (n=24) (D). (E) Plasma cholesterol levels were numerical increased, though not statistically significant, in *Cre*-negative *Ccl2*^flx/flx^ (n=15, black squares) compared with *Ccl2*^wt/wt^ (n=17, grey triangles) mice 11 weeks after rAAV-PCSK9 injection. Graphs show mean ± SEM. \**p*<0.05; \*\**p*<0.01; NS, not statistically significant in a mixed-effects model for repeated measurements with Šidák’s multiple comparison test (A,D,E) and Welch t-test (C).

Sterol analysis was performed to provide clues to the source of the increased plasma cholesterol in *Ccl2*^SMC-KO^ mice **(Fig. 3C)** [29]. Markers for the rate of cholesterol synthesis (lathosterol and desmosterol) and intestinal cholesterol absorption (campesterol, stigmasterol, sitosterol, lanosterol) were either unchanged or reduced in plasma of *Ccl2*^SMC-KO^ compared with *Ccl2*^SMC-WT^ mice. These observations instead pointed towards a potential difference in LDL clearance, which could be caused by a difference in rAAV8 transduction efficiencies leading to differences in PCSK9 levels. We therefore measured plasma PCSK9 levels in samples taken at 2 and 11 weeks after rAAV8. Plasma PCSK9 levels increased from 2 to 11 weeks, but no statistically significant differences were found between *Ccl2*^SMC-KO^ and *Ccl2*^SMC-WT^ mice (**Fig. 3D**). Furthermore, introducing plasma PCSK9 levels in multiple regression analysis did not weaken the link between genotype and plasma cholesterol levels (**Supplemental Table 1**).

Although the floxed *Ccl2* mice were already backcrossed into the B6 mouse background 10 times when we acquired them and further 3-4 times in our facility, a region around the *Ccl2* locus will be derived from the 129/SVJ embryonic stem cells in which the *Ccl2* gene targeting was performed [26]. This passenger gene region could potentially harbor plasma cholesterol regulating loci. Furthermore, the genetic modification of the *Ccl2* locus could affect neighboring gene expression in any cell type, which in theory could impact lipoprotein metabolism. To explore this, we also analyzed plasma lipids in CreER^T2^-negative *Ccl2*^flx/flx^ and *Ccl2*^wt/wt^ mice that had been subjected to a similar regimen of rAAV8-PCSK9 treatment and high-fat diet. We found a numerical increase in plasma cholesterol levels in *Ccl2*^flx/flx^ compared with *Ccl2*^wt/wt^ at the last time point (**Fig. 3E**). Although this difference was not significant (*p*=0.08), it is consistent with genetic background effects, rather than the SMC-specific *Ccl2* deletion, being the underlying cause of the differences in plasma cholesterol.

### Atherosclerosis development

Atherosclerosis development was assessed in sections of the aortic root and by en face analysis of the aortic arch after 12 weeks of hypercholesterolemia (**Fig. 4A-B**). Aortic root plaque area was significantly increased in *Ccl2*^SMC-KO^ compared with *Ccl2*^SMC-WT^ mice (**Fig. 4C**). In the aortic arch, the lesion coverage showed an insignificant tendency towards an increase in *Ccl2*^SMC-KO^ compared with wildtype littermates (**Fig. 4D**).

**Figure 4.**
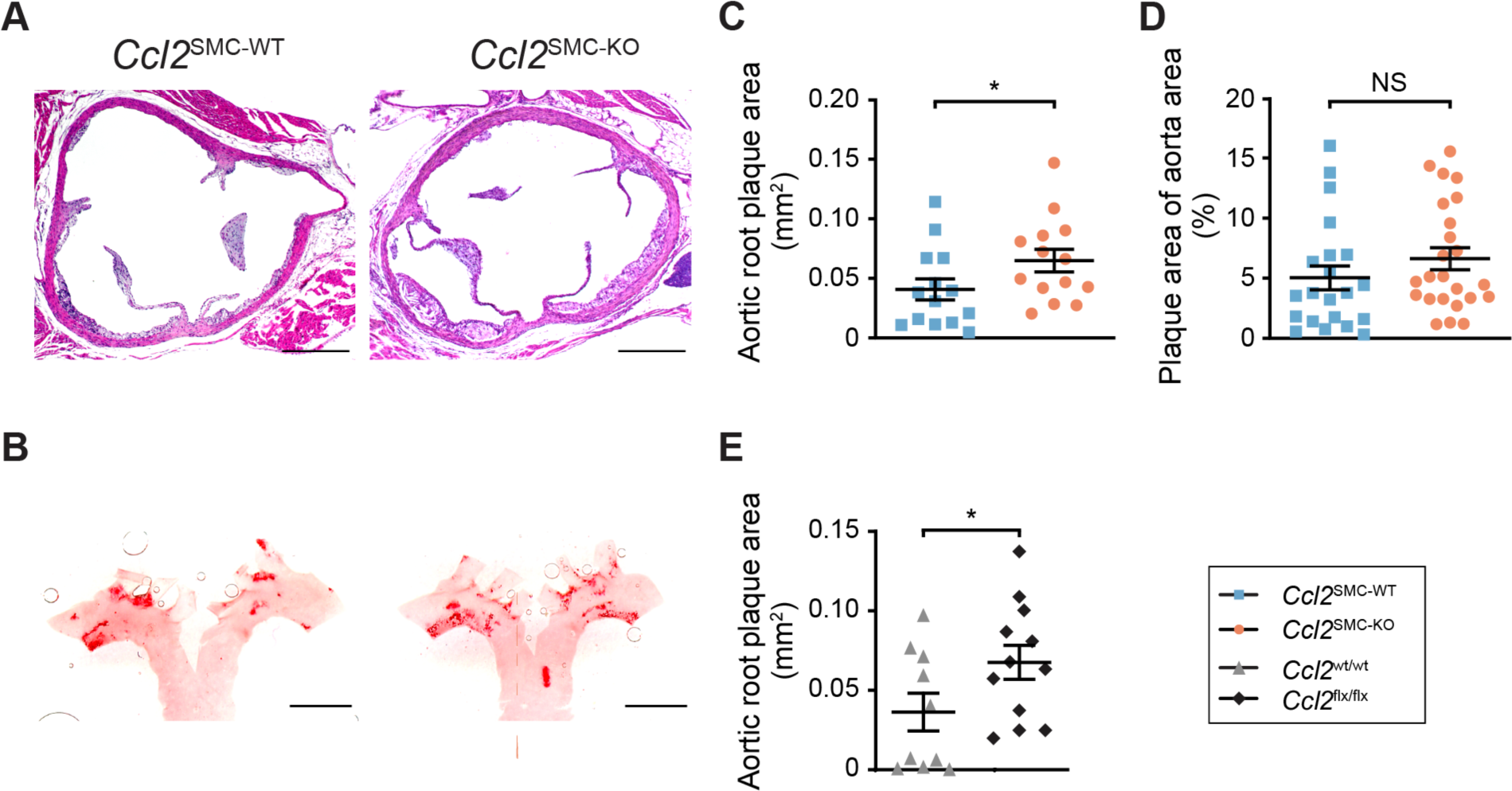
Quantification of atherosclerosis after 12 weeks of atherogenic diet feeding. Representative images of (A) hematoxylin-eosin-stained cross sections from the aortic root, and (B) Oil Red O-stained aortic arches opened for en face quantification. Scale bars = 400μm (A) and 2 mm (B). (C) Plaque size was statistically significantly increased in *Ccl2*^SMC-KO^ (n=14, orange circles) mice compared with *Ccl2*^SMC-WT^ (n=14, blue squares) mice in the aortic root. (D) In the aortic arch, a similar tendency was seen but not statistically significant (*Ccl2*^SMC-KO^, n=24; *Ccl2*^SMC-WT^, n=21). Aortic root plaque size was also statistically significantly increased in *Cre*-negative *Ccl2*^flx/flx^ (black squares, n=12) mice compared with *Ccl2*^wt/wt^ (grey triangles, n=10) (E). Graphs show mean ± SEM. * *p*<0.05, NS, not statistically significant by Welch t-test on log-transformed data.

Aortic root cross-sections were analyzed for plaque cellular composition using the macrophage marker LGALS3 and the SMC-marker ACTA2 (**Fig. 5A**). LGALS3 can be expressed on SMC lineage cells in addition to macrophages. Still, we have previously found that the level of misclassification by standard immune fluorescent staining in PCSK9-induced atherosclerosis is minor as assessed by SMC lineage tracing [30]. The number of LGALS3+ cells was increased in *Ccl2*^SMC-KO^ compared with wildtype littermates (**Fig. 5B**), while the number of ACTA2+ cells was unaffected (**Fig. 5C**).

**Figure 5.**
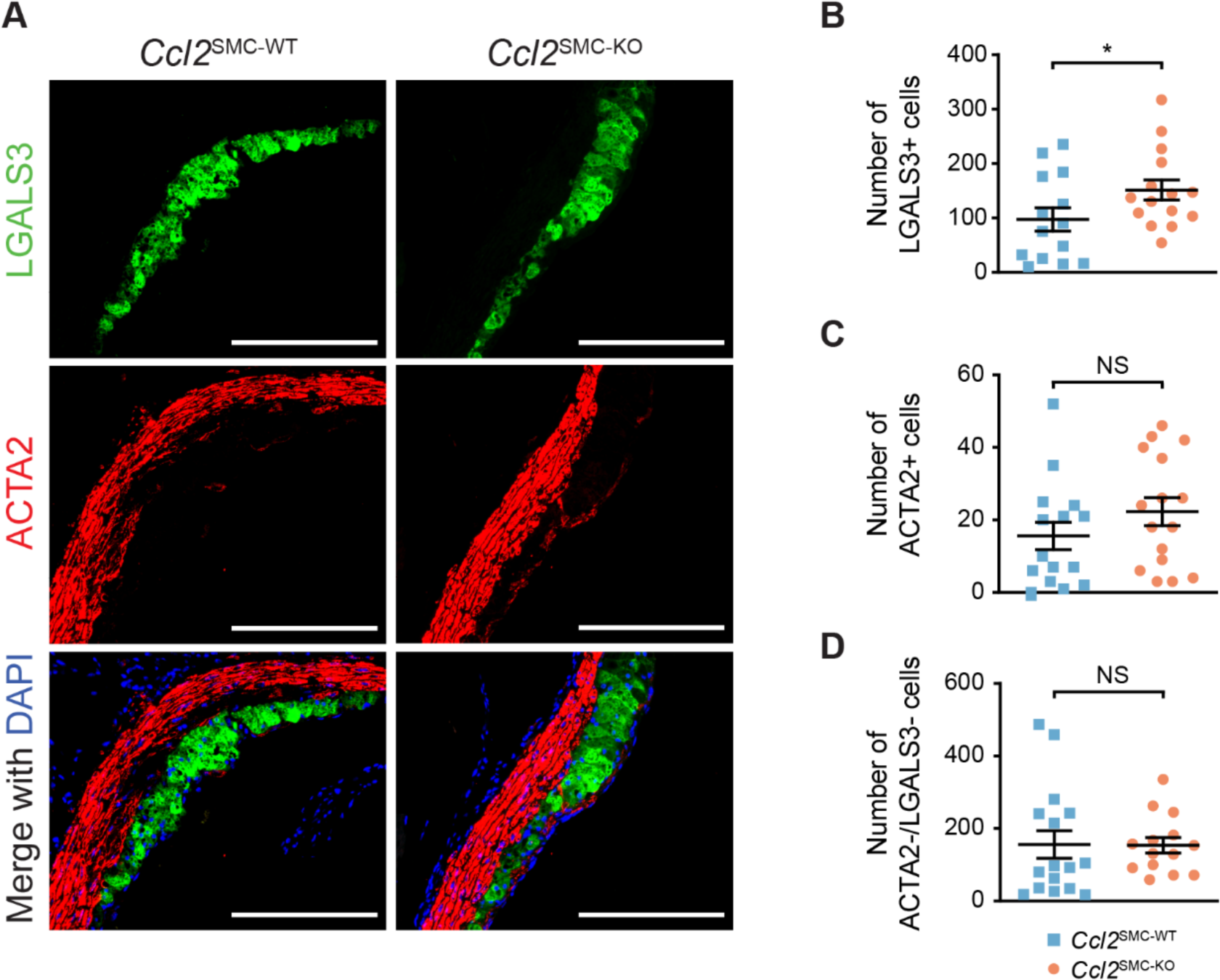
Plaque composition in the aortic root after 12 weeks of atherosclerosis development. (A) Representative images showing LGALS3+ cells (green), ACTA2+ cells (red), and merged channels with DAPI (blue) from *Ccl2*^SMC-WT^ and *Ccl2*^SMC-KO^ mice. Scale bars 200 μm. Number of plaque LGALS3+ cells (*Ccl2*^SMC-WT^, n=14; *Ccl2*^SMC-KO^, n=15) (B), number of ACTA2+ cells (*Ccl2*^SMC-WT^, n=15; *Ccl2*^SMC-KO^, n=16) (C) and number of ACTA2/LGALS3 double-negative cells (*Ccl2*^SMC-WT^, n=16; *Ccl2*^SMC-KO^, n=14) (D) in aortic root sections from *Ccl2*^SMC-WT^ (blue squares) and *Ccl2*^SMC-KO^ (orange circles). Graphs show mean ± SEM. \**p*<0.05; NS, not statistically significant by Welch t-test on log-transformed data.

Murine atherosclerotic lesions also contain a large population of SMC lineage cells that lose expression of ACTA2 and modulate to other forms [31]. We therefore also quantified ACTA2/LGALS3 double-negative cells (ACTA2-/LGALS3-), which are enriched for modulated plaque SMCs, but found no differences between the two groups (**Fig. 5D**).

None of the differences in atherosclerosis measures between groups were statistically significant in a multiple regression model incorporating final plasma cholesterol (data not shown) or the area-under-the-curve of plasma cholesterol levels from 2 until 11 weeks (**Supplemental Tables 2** and **3**). This indicates that the observed changes were explained by the unexpected differences in plasma cholesterol between groups, which was unrelated to the deletion of Ccl2 in SMCs. This was confirmed by analysis of aortic root plaque area in CreER^T2^-negative *Ccl2*^flx/flx^ and *Ccl2*^wt/wt^ mice (**Fig. 4E**). As observed for the *Ccl2*^SMC-KO^ mice, the plaque area was significantly increased in *Ccl2*^flx/flx^ mice compared to their wildtype counterparts, supporting the conclusion that secretion of CCL2 by modulated SMCs in the plaque is not an important mediator in plaque development.

## Discussion

SMCs have traditionally been considered to have a protective role in atherosclerosis by producing fibrous tissue that stabilizes the plaque and protects it from rupture and thrombotic complications [32]. Over the last decade, however, multiple studies have revisited the role of SMCs in atherosclerosis using mouse models and lineage-tracing techniques [31]. Such experiments have shown that plaque SMCs derive from a small group of arterial SMCs that undergo clonal expansion and modulate to cells with fibroblast-like (aka. fibromyocytes) and chondrocyte-like (aka. chondromyocytes) phenotypes [33–38]. Some of these show evidence of inflammatory signaling [39], but the causal role of this for plaque development is still sparsely understood.

In the present study, we tested whether the ability of SMCs to secrete the proinflammatory cytokine CCL2 is a mediator linking SMC modulation to arterial inflammation and plaque development. CCL2 is a central cytokine involved in monocyte recruitment to tissues, and global knockout of *Ccl2* reduces plaque formation and progression in mice [6–8,13]. We confirmed the expression of CCL2 in human plaques in SMC-rich areas including the arterial media in the pre-diseased arterial wall. Furthermore, we confirmed the ability of both human and mouse SMCs to respond to an inflammatory stimulus with CCL2 expression. To test the potential causal role of this, we studied atherosclerosis in mice with SMC-specific conditional knockout of *Ccl2*.

Unexpectedly, we found increased hypercholesterolemia and atherosclerosis in mice with SMC-specific deficiency of *Ccl2* compared with wildtypes littermates. Importantly, however, a critical control experiment in mice lacking the Cre recombinase showed that this paradoxical effect was not caused by the knockout of *Ccl2* in SMCs. Rather it was explained by genetic differences between the floxed and wildtype *Ccl2* allele; either the floxing of the gene itself or passenger gene variants in linkage disequilibrium with the *Ccl2* locus originating from the 129/SVJ embryonic stem cells used to create the mouse line [22,40]. The genetic differences appeared to accelerate atherosclerosis at least partly by increasing hypercholesterolemia. Plasma cholesterol was significantly increased in *Ccl2*^SMC-KO^ compared with *Ccl2*^SMC-WT^ mice, which was unexpected since previous studies of mice with global CCL2 knockout or overexpression did not show any effects on plasma cholesterol levels [6–8,13]. Although the differences in plasma cholesterol levels in *Ccl2*^flx/flx^ and *Ccl2*^wt/wt^ mice (without the *Cre* transgene) were not statistically significant, they resembled those of the main study. The differences were not caused by differences in the efficiency of rAAV-PCSK9 transduction since the effect was not present after 2 weeks but developed over time and plasma PCSK9 levels were similar between groups in the two genotypes.

While we were finalizing our studies, Owsiany et al. reported increased atherosclerosis and macrophage recruitment in *Apoe*^-/-^ mice with heterozygous, but not homozygous, knockout of *Ccl2* in SMCs [20]. The increased atherosclerosis in heterozygous animals appeared to result from increased monocytosis in combination with a preserved ability to recruit monocytes into the arterial intima, which was lost in homozygous animals [20]. Recombination rates were not quantified in the study of Owsiany et al. and hence it is difficult to compare our mice (with an app. 75% recombination rate of the floxed *Ccl2* locus) with the two genotypes reported in that study. Notably, even though our findings may on the surface seem consistent with the increased atherosclerosis reported in the heterozygous mice in Owsiany et al., the proatherogenic effect in our experiments, as discussed above, was not caused by *Ccl2* deletion. It is unlikely that the genetic background effects we saw in our study influenced the results of Owsiany et al., because they used a different floxed *Ccl2* mouse line, which was created using C57BL/6J embryonic stem cells [20,41].

Overall, we conclude that the knockout of *Ccl2* in SMCs in our experimental context did not change atherogenesis, similar to the homozygous SMC knockout in Owsiany et al. [20]. The most likely reason is that SMC-secreted CCL2 is not an important regulator of plaque inflammation. It should be noted, however, that upregulation of both *Ccl7* (*Mcp3)* and *Ccl12* (*Mcp5)* located downstream of the *Ccl2* gene was originally demonstrated for the mouse line used in our studies upon Cre-mediated deletion of the floxed *Ccl2*-region [22]. Given that both encoded proteins are also ligands for the C-C chemokine receptor type 2 (CCR2), potential compensating effects from these genes cannot be excluded.

### Strengths and limitations

Our study has limitations. Recombination of CCL2 alleles in SMCs was not complete as is generally the case in conditional knockout studies. This raises the question whether more robust inhibition of CCL2 secretion from SMCs would have had an effect on plaque development. It is also possible that remaining CCL2-competent SMCs could outcompete recombined cells during clonal expansion in plaques if CCL2 secretion confers a survival benefit. Finally, our studies were not designed to investigate the impact of SMC-specific knockout on monocyte trafficking, which would have been relevant on the background of the recent findings by Owsiany et al. [20].

## Conclusion

Conditional deletion of *Ccl2* in SMCs does not inhibit the development of atherosclerosis in mice with PCSK9-induced hypercholesterolemia.

## Author contribution

Participated in research design: SG, JFB

Performed experiments and data analysis: SG, FL, UJFT, JTS, CBS

Writing or contributing to the writing of the manuscript: SG, UJFT, CBS, JFB

Review and final approval of the manuscript: SG, FL, UJFT, JTS, CBS, JFB

## Acknowledgements

We thank Lisa Maria Røge and Dorte Qualmann for excellent technical assistance.

## Abbreviations and Acronyms

ACTA2: smooth muscle alpha-2 actin
CCL2: C-C motif chemokine ligand 2
CCR2: C-C chemokine receptor type 2
EC: endothelial cell
FCS: fetal calf serum
FITC: fluorescein isothiocyanate
FPLC: fast protein liquid chromatography
hVSMC: human aortic smooth muscle cell
IKKβ: inhibitor of nuclear factor kappa-B kinase subunit beta
IL-1β: interleukin-1 beta
IL-1R: interleukin-1 receptor
LGALS3: galectin 3
MOVAS: mouse vascular smooth muscle cell line
MYH11: myosin heavy chain 11
NF-κB: nuclear factor kappa-light-chain-enhancer of activated B cells
PBS: phosphate-buffered saline
PCSK9: proprotein convertase subtilisin/kexin type 9
PFA: phosphate-buffered formaldehyde
rAAV: recombinant adeno-associated virus
SMC: smooth muscle cell

