## Supplemental Material for "Conditional deletion of *Ccl2* in smooth muscle cells does not reduce early atherosclerosis in mice"

[a]Department of Clinical Medicine, Aarhus University, Aarhus, Denmark.
[b]Department of Cardiology, Aarhus University Hospital, Aarhus, Denmark.
[c]Division of Clinical Chemistry, Department of Laboratory Medicine, Karolinska Institutet, Stockholm, Sweden.
[d]Clinical Chemistry, Karolinska University Laboratory, Karolinska University Hospital, Stockholm, Sweden.
[e]Centro Nacional de Investigaciones Cardiovasculares Carlos III (CNIC), Madrid, Spain.

### Supplemental Methods

#### Genotyping

Mice were genotyped by PCR using primers (5'-AAGACTCCGCTCAGCCACT-3' and 5'-GAGCCAACTGGAATGTCTGG-3') flanking the loxP site upstream of exon 1 in the *Ccl2* gene.

#### Recombination efficiency

Pieces of aorta were isolated from *Myh11*-CreER$^{T2}$ x *Ccl2*$^{flx/flx}$ and *Ccl2*$^{flx/flx}$ mice. The adventitial layer was removed and the endothelial cells (ECs) scraped off with a scalpel. DNA was extracted from the remaining arterial medial layer using the QIAamp DNA FFPE Tissue Kit (Qiagen, cat. no. 56404). Quantitative PCR for the non-recombined *Ccl2*$^{flx}$ allele was performed with 7.5 ng genomic DNA serving as a template for PCR amplification using Maxima SYBR Green/ROX qPCR Mastermix (Thermo Fischer Scientific cat. no. K0221) and *Ccl2*-specific primers (5'- AGATACCTGAGTGGAAGACTCC-3' and 5'-ACTACTCCTGGTAGCTCTCT-3') with the latter primer located within the region flanked by loxP sites. Mouse beta-actin (*Actb)* served as reference gene and was amplified using the primers 5'-CAGCCAACTTTACGCCTAGC-3' and 5'-TCTCAAGATGGACCTAATACGG-3'. The qPCR protocol for both amplicons consisted of an initial polymerase activation/denaturation step of 10 minutes at 95˚C followed by 40 cycles of 30 seconds at 95˚C, 1 minute at annealing temperature, 1 minute at 72˚C and last, 1 cycle of 1 minute at 95˚C, 30 seconds at 60˚C and 30 seconds at 95˚C. Annealing temperatures were 66°C and 64°C for *Ccl2*$^{flx}$ primers and *Actb* primers, respectively. Average recombination efficiency was determined as the average reduction in *Ccl2*$^{flx}$ allele levels (normalized to *Actb*) in genomic DNA from *Myh11*-CreER$^{T2}$ x *Ccl2*$^{flx/flx}$ compared with *Ccl2*$^{flx/flx}$ mice and calculated by the delta-delta Ct method.

#### Blood analysis

Total plasma cholesterol levels (non-fasting) were measured in duplicates with an enzymatic reagent (CH201, Randox). Distribution of cholesterol over the different lipoprotein subclasses was determined in pooled plasma samples from the respective experimental groups by fast protein liquid chromatography (FPLC, Äkta Purifier system, GE Health,

Uppsala, Sweden) using a Superose 6 column and a flow rate of 0.5 ml/min essentially as described [1]. Individual fractions were assayed for cholesterol content using commercially available reagents (Roche Diagnostic, Basel, Switzerland). Plasma sterol analyses were performed by gas-liquid chromatography using an established protocol as detailed previously [1].

**Flow cytometry**

MOVAS cells were seeded with a density of $6.25x10^3$ cells/$cm^2$ in 12-well plates. After 24 hours, cells were incubated with IL-1β 1 ng/ml for 18 hours. Brefeldin A (Sigma-Aldrich, cat. no. B7651,10 μg/ml) was added in the last 8 hours of incubation to inhibit protein transport and increase intracellular staining signal. The media was then removed, and the cells were trypsinized, pelleted, and washed with Staining Buffer (PBS with 1% FCS, 0.09% $NaN_3$, pH 7.5). Permeabilization and fixation were done using the Fixation/Permeabilization Solution Kit (BD Bioscience cat no. 554714). Cells were stained for 30 min at 4˚C with a fluorescein isothiocyanate (FITC)-conjugated hamster (Armenian) antibody against CCL2 (LifeSpan Biosciences, cat. no. LS-C105897, 1:50). A FITC-conjugated isotype control was used to identify unspecific staining (LifeSpan Biosciences, cat. no. LS-C292320). All samples were run on a 3-laser NovoCyte 1116-171 Analyzer (NovoCyte) and data analyzed using FlowJo.

**Supplemental Tables**

| Regression model<br><br>$Y = \boldsymbol{\beta_1} \cdot B + \boldsymbol{\beta_2} \cdot \mathrm{C} + \boldsymbol{\beta_0}$<br><br>where $Y$ is plasma total cholesterol at 11 weeks (in mM), B is a binary variable representing genotype and C is plasma PCSK9 levels at 11 weeks (in µg/mL). | | |
|---|---|---|
| | Estimate | P value |
| $\beta_0$ | 7.85 (95%CI: 3.12 -12.57) | 0.0017 |
| $\beta_1$ | 5.88 (95%CI: 2.88 – 8.89) | **0.0003** |
| $\beta_2$ | 0.25 (95%CI: 0.15 – 0.36) | **<0.0001** |

**Supplemental Table 1. Determinants of plasma total cholesterol.** Multiple regression analysis to analyze the independent effect of genotype and plasma PCSK9 on plasma cholesterol levels.

| Regression model<br><br>$Y = \boldsymbol{\beta_1} \cdot B + \boldsymbol{\beta_2} \cdot \mathrm{C} + \boldsymbol{\beta_0}$<br><br>where *Y* is log-transformed plaque area (in $mm^2$), B is a binary variable representing genotype and C is area-under-the-curve for plasma total cholesterol (in mM·week). | | |
|---|---|---|
| | Estimate | P value |
| $\beta_0$ | -2.10 (95%CI: -2.97 – 1.22) | <0.0001 |
| $\beta_1$ | -0.11 (95%CI:-0.46 – 0.24) | 0.5245 |
| $\beta_2$ | $4.10 \cdot 10^{-3}$ (95%CI:$0.066 \cdot 10^{-3}$ – $8.13 \cdot 10^{-3}$) | **0.0466** |

**Supplemental Table 2. Determinants of plaque area in *Ccl2*$^{\text{SMC-KO}}$ and *Ccl2*$^{\text{SMC-WT}}$ mice.** Multiple regression analysis to analyze the independent effect of genotype and area under the curve for plasma total cholesterol on plaque area.

| Regression model<br><br>$Y = \boldsymbol{\beta_1} \cdot B + \boldsymbol{\beta_2} \cdot \mathrm{C} + \boldsymbol{\beta_0}$<br><br>where *Y* is log-transformed LGALS3+ area (in $mm^2$), B is a binary variable representing genotype and C is area-under-the-curve for plasma total cholesterol (in mM·week). | | |
|---|---|---|
| | Estimate | P value |
| $\beta_0$ | 0.91 (95%CI:0.11 – 1.71) | 0.0275 |
| $\beta_1$ | -0.06 (95%CI:-0.39 – 0.27) | 0.7096 |
| $\beta_2$ | $5.72 \cdot 10^{-3}$ (95%CI: $2.06 \cdot 10^{-3}$ – $9.39 \cdot 10^{-3}$) | **0.0037** |

**Supplemental Table 3. Determinants of plaque LGALS3+ cells in *Ccl2*$^{SMC\text{-}KO}$ and *Ccl2*$^{SMC\text{-}WT}$ mice.** Multiple regression analysis to analyze the independent effect of genotype and area under the curve for plasma total cholesterol on LGALS3+ area.
